# Evaluation of Oxford Nanopore MinION^TM^ Sequencing for 16S rRNA Microbiome Characterization

**DOI:** 10.1101/099960

**Authors:** Xiao Ma, Elyse Stachler, Kyle Bibby

## Abstract

In this manuscript we evaluate the potential for microbiome characterization by sequencing of near-full length 16S rRNA gene region fragments using the Oxford Nanopore MinION (hereafter ‘Nanopore’) sequencing platform. We analyzed pure-culture *E. coli* and *P. fluorescens*, as well as a low-diversity mixed community sample from hydraulic fracturing produced water. Both closed and open reference operational taxonomic unit (OTU) picking failed, necessitating the direct use of sequences without OTU picking. The Ribosomal Database Project classifier against the Green Genes database was found to be the optimal annotation approach, with average pure-culture annotation accuracies of 93.8% and 82.0% at the phyla and genus levels, respectively. Comparative analysis of an environmental sample using Nanopore and Illumina MiSeq sequencing identified high taxonomic similarity when using a weighted metric (Bray-Curtis), and significantly reduced similarity when using an unweighted metric (Jaccard). These results highlight the great potential of Nanopore sequencing to analyze broad microbial community trends, and the challenge of applying Nanopore sequencing to discern rare taxa in mixed microbial communities. Finally, we observed that between-run carryover following washes on the same flowcell accounted for >10% of sequence reads, necessitating future development to either prevent carryover or filter sequences of interest (e.g. barcoding).

## Introduction

Interest in studying the microbiome, microbiota associated with various environments, has exploded in recent years largely due to the rapid expansion in ‘next-generation sequencing’ capabilities and subsequent reduction in cost. It has been recognized that the human microbiome plays an important role in many different clinical outcomes, including obesity (1), immune state (2), and infection (3). The human microbiome is comprised of a diverse array of commensal microorganisms in and on the human body, and emerging research has suggested a clinical role for the microbiome in either therapeutic development (e.g. probiotics) (4) or diagnostics (5). Concurrently, significant interest has emerged in the microbiomes of various other environments, such as buildings (6-8) and water systems (9).

Currently, the most common microbiome analysis approach is high-throughput amplicon sequencing of the 16S rRNA gene region. The most widely used sequencing technologies are Illumina sequencing platforms (e.g. the MiSeq and HiSeq). These platforms are accurate and generate a large amount of data, but are limited by capital costs, the necessity to pool samples to reduce per-sample costs, sequence read length, and a turnaround time of days to weeks that may be inadequate for many applications. Recently, Oxford Nanopore has released a small and inexpensive sequencing platform called the MinION^TM^, which has been previously reviewed (10, 11). Capital costs are reduced to per-run costs that are comparable with the current Illumina platforms, and data analysis is possible in near-real time, enabling investigators to generate sequence data as-needed. For example, samples were correctly assigned to the *Salmonella* species in 20 minutes and serotype in 40 minutes (12). Amplicon (non-metagenomic) sequencing has also been successfully employed to identify both bacterial and viral origin (13, 14). Additionally, this technology routinely produces sequences >10kb in length, enabling sequencing of the full 16S rRNA gene region and more reliable taxonomic placement. Despite the benefits, the primary drawback to this technology is relatively high error rates (currently reported to be ~8%), hindering metagenomic analysis of highly diverse microbiome samples and requiring additional development and validation. Previous investigations have demonstrated successful 16S rRNA microbial community classification via sequencing of a single mouse gut microbiome (15) and a single mock microbial community (16), including species-level assignment (16); however, a formal assessment of 16S rRNA sequencing on the Nanopore platform, including analysis of pure-culture samples for annotation validation, is currently lacking. Development of Nanopore technology for microbiome analysis would enable rapid (<12 hours from sample to results) and low cost microbiome characterization that would be applicable to both the clinic and other applications.

In the current study we evaluate the potential for microbiome characterization via Nanopore sequencing of near full-length 16S rRNA PCR amplicons. First, we evaluate the accuracy and annotation strategies of sequences from pure culture *E. coli* and *P. fluorescens* to determine the most appropriate sequence analysis approach. We then compare performance of Nanopore sequencing against the current state of the art Illumina sequencing using a sample from hydraulic fracturing wastewater. Finally, we investigate an apparent between-run carryover phenomena, and propose necessary future investigations to enable 16S rRNA microbiome characterization on the Nanopore platform.

## Results and Discussion

### Sequencing

Sequencing was performed on eight 16S rRNA libraries; however, due to apparent between-run carryover (discussed below), only the initial run on each of three flow cells was used for subsequent analyses of annotation approach and accuracy. These sequencing libraries were derived from two pure-culture samples, *E. coli* and *P. fluorescens*, and a low-diversity sample from hydraulic fracturing produced water. Detailed sequencing results are shown in Table 1. Raw sequence data can be found on Figshare using DOI: https://dx.doi.org/10.6084/m9.figshare.4515752.v3.

**Table 1.**
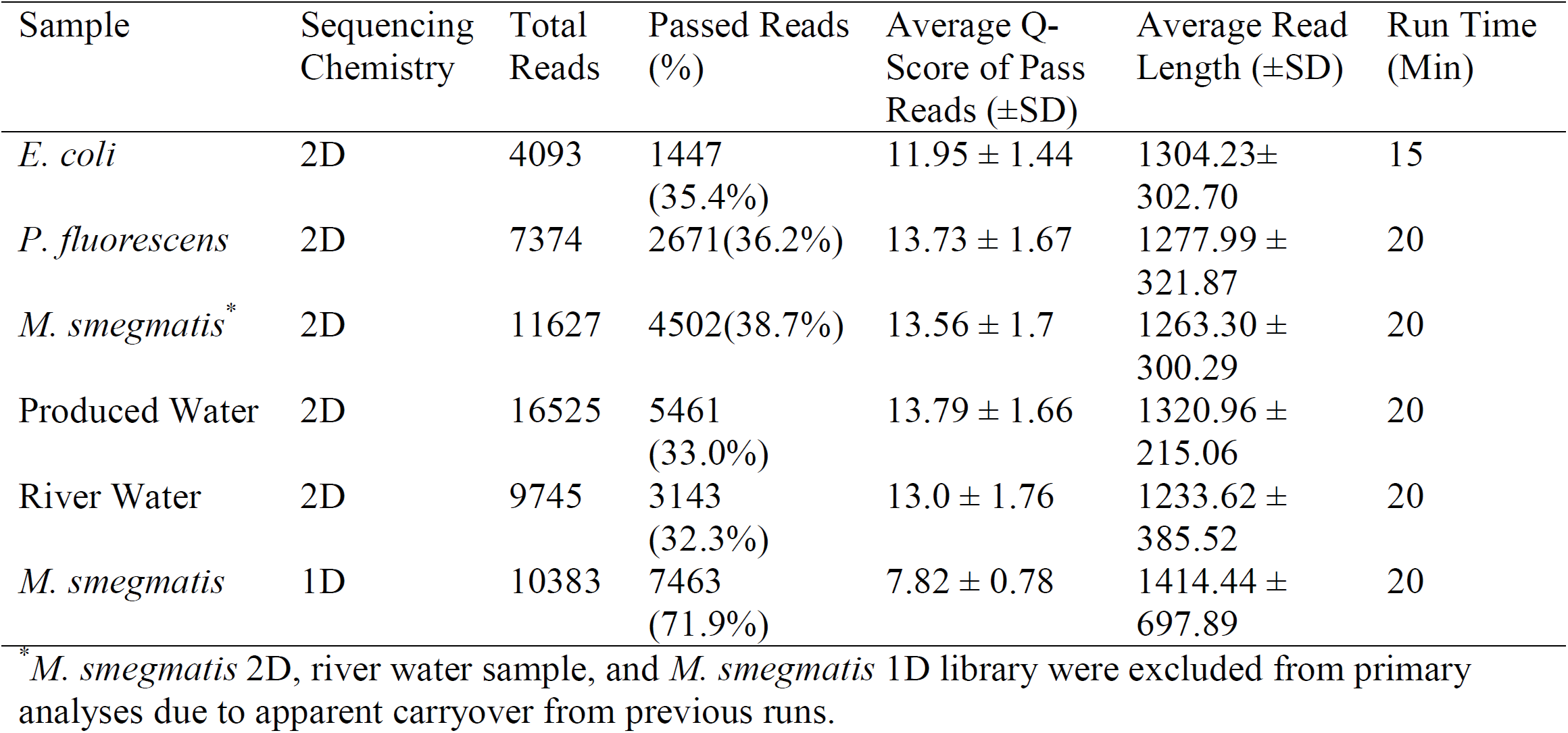
Sequencing results and output from Oxford Nanopore MinION runs.

### Pure Culture Analyses

We first evaluated the suitability of Operational Taxonomic Unit (OTU) picking methods using Nanopore sequence data. OTU picking via comparison with a reference library, i.e. closed-reference OTU picking, failed, with no sequences being assigned to an OTU (i.e. all sequences were excluded). *De novo* OTU clustering, i.e. OTU clustering by determining between-sequence similarity, was then evaluated using similarity thresholds between 90-100%. At the typically used similarity threshold of 97%, all sequences from both pure-culture samples were assigned to unique OTUs (i.e. the ratio of OTUs to sequences was one). Results from *de novo* OTU clustering evaluation are shown in Figure 1. These results highlighted the challenge of clustering reads from long, error-prone sequences, and necessitated analyzing the taxonomy of sequences individually without OTU picking.

**Figure 1.**
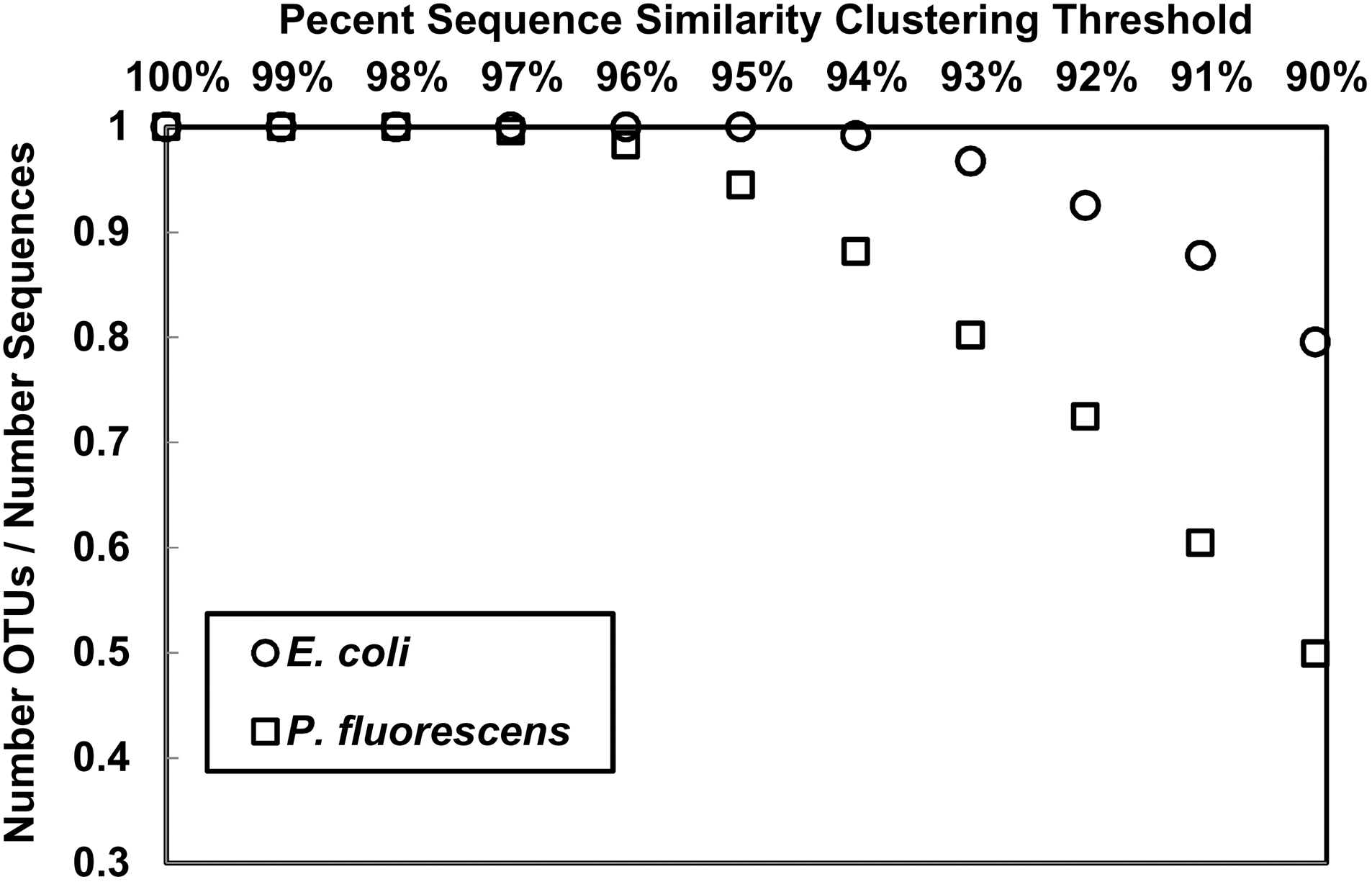
Number of observed *de novo* operational taxonomic units (OTUs) per number of sequences at different similarity thresholds for *E. coli* and *P. fluorescens* pure culture DNA sample sequenced by Nanopore.

We next evaluated the ability to accurately annotate the taxonomy of pure-culture Nanopore 16S rRNA sequences using three different annotation approaches: the naïve Bayesian Ribosomal Database Project (RDP) classifier with the Green Genes database; the RDP classifier against the RDP database; and BLAST against the Green Genes database. Results from this evaluation are shown in Figure 2. The RDP classifier against the Green Genes database was found to be the best performing annotation strategy. Using this approach, the annotation accuracy for *E. coli* was 96.7% and 81.9% at the phyla and genus level, respectively, and was 90.9% and 82.0% for *P. fluorescens* at the phyla and genus level, respectively.

**Figure 2.**
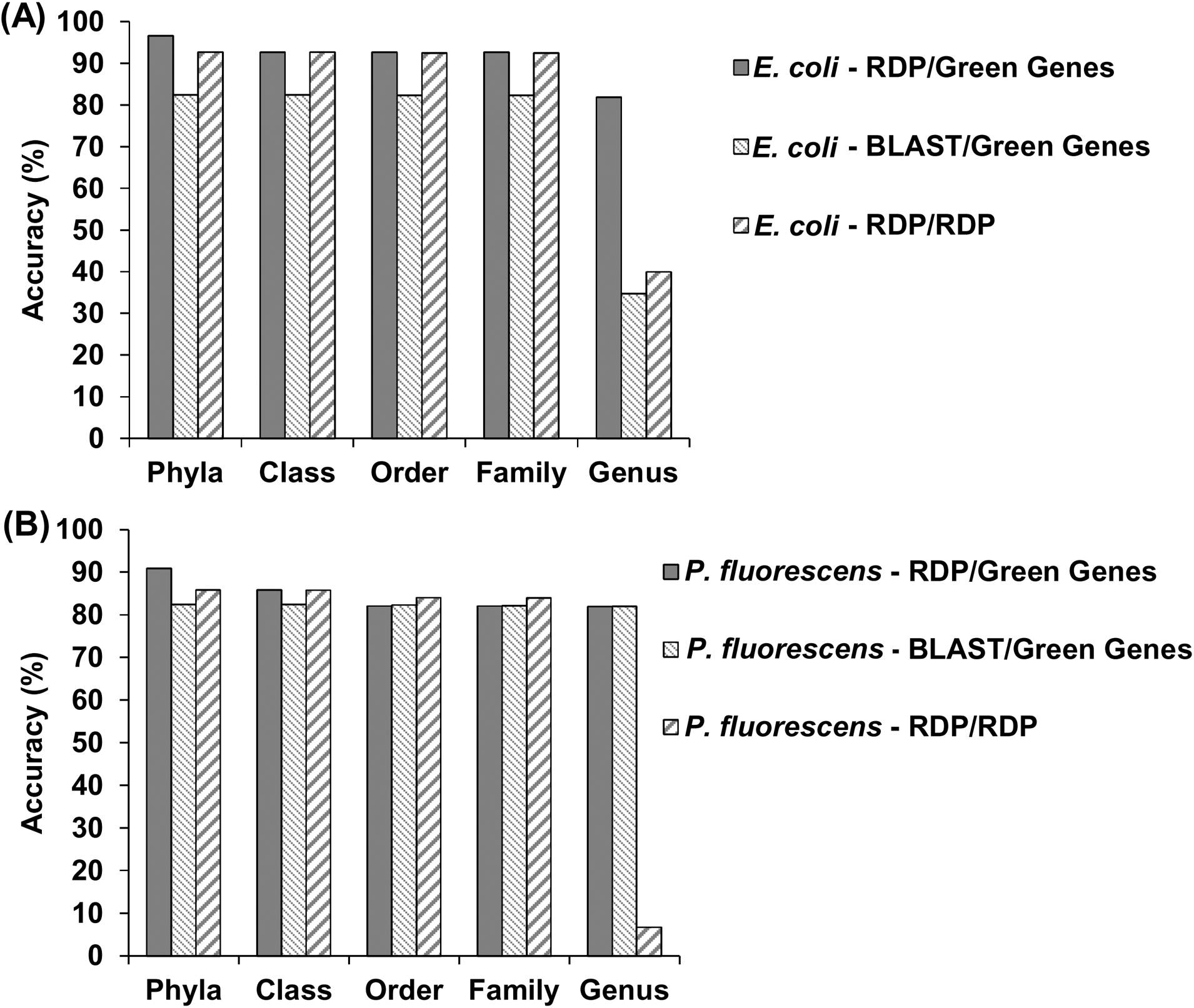
Accuracy of taxonomy assignment at different phylogenetic levels for (A) *E. coli*; and (B) *P. fluorescens*

### Comparison with Illumina Sequencing

We next evaluated Nanopore sequencing to characterize an environmental sample from hydraulic fracturing produced water (17). This sample was selected as it exhibited low alpha-diversity in previous analyses. The same DNA extract was used for both analyses.

By assigning taxonomy to each individual sequence, five phyla were identified by the Nanopore platform, and eleven phyla were identified by the Illumina platform. Among them, four phyla that together accounted for greater than 99% of sequence relative abundance were identified by both platforms (Table 2). Nine shared genera were detected by both platforms, accounting for relative abundances of 98.3% on the Nanopore platform and 81.6% on the Illumina platform (Table 2). Both the Nanopore and Illumina platforms revealed similar microbial community structure of the produced water sample. The Firmicutes Phylum dominated the microbial community with relative abundance higher than 90% with both platforms (Table S1). Phyla unique to the Nanopore and Illumina platforms accounted for less than 0.5% of relative abundance (Table S1). At the genus level, the produced water microbial community was dominated by the genus *Halanaerobium* (Table S2), with 95.6% of Nanopore sequences and 76.8% of Illumina sequences being assigned to *Halanaerobium*. In addition, 14.6% of Illumina 16S rRNA sequence reads were assigned to Clostridiales, which is within the same class with *Halanaerobium* (Clostridia) (Table S2).

**Table 2.**
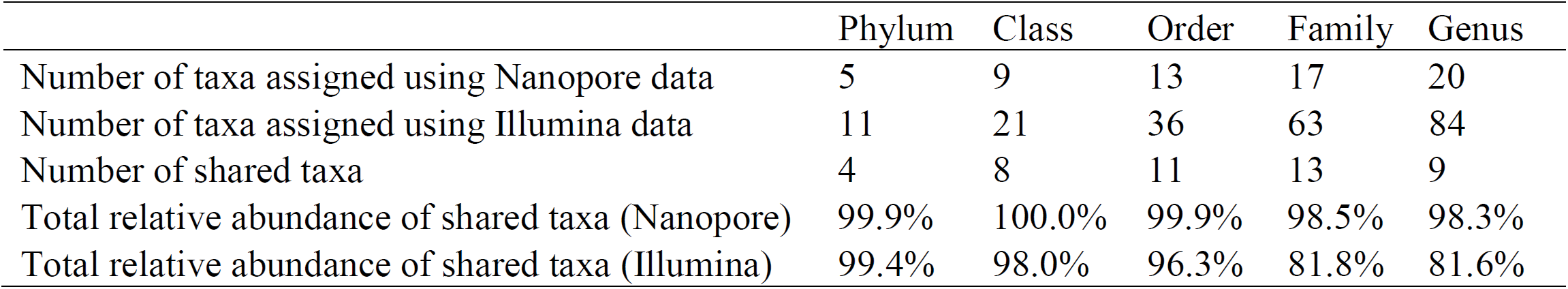
Comparison of number of taxa at different phylogenetic levels for the produced water sample

|  | Phylum | Class | Order | Family | Genus |
| --- | --- | --- | --- | --- | --- |
| Number of taxa assigned using Nanopore data | 5 | 9 | 13 | 17 | 20 |
| Number of taxa assigned using Illumina data | 11 | 21 | 36 | 63 | 84 |
| Number of shared taxa | 4 | 8 | 11 | 13 | 9 |
| Total relative abundance of shared taxa (Nanopore) | 99.9% | 100.0% | 99.9% | 98.5% | 98.3% |
| Total relative abundance of shared taxa (Illumina) | 99.4% | 98.0% | 96.3% | 81.8% | 81.6% |

We also calculated the Jaccard distance (solely based on presence/absence of each taxa) and Bray-Curtis distance (based on both presence/absence and relative abundance of each taxa) between the produced water microbial community revealed by Nanopore and Illumina platforms at different taxonomic levels. Jaccard and Bray-Curtis distances measure the degree of microbial community structure difference between two samples, with a value of one indicating no community structure overlap and a value of zero indicating identical microbial communities. We adopted these measures to evaluate the level of difference between the technical replicates of the same produced water sample between the Nanopore and Illumina sequencing platforms. The Jaccard distance increased from 0.62 at the phylum level to 0.89 at genus level; the Bray-Curtis distance increased from 0.04 at phylum level to 0.22 at genus level (Figure 3). Jaccard distances were higher than Bray-Curtis distances at all phylogenetic levels, because more taxa were assigned using short sequence reads by Illumina sequencing (Table 2) and Jaccard distance only accounts for the presence and absence of each assigned taxa whereas Bray-Curtis distances take relative abundance into account.

**Figure 3.**
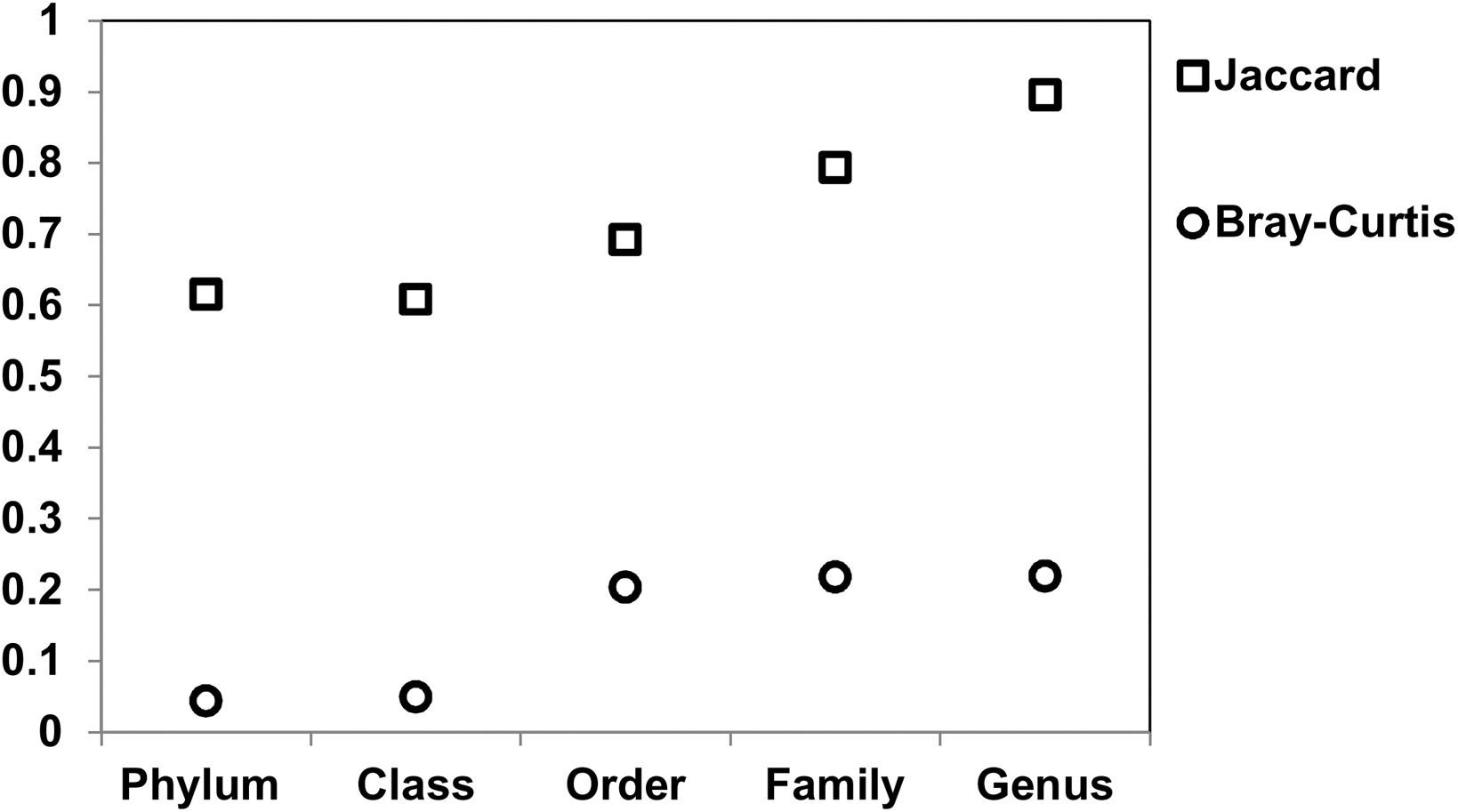
Jaccard (unweighted) and Bray-Curtis (weighted) dissimilarity values for a hydraulic fracturing produced water sample analyzed by Nanopore and Illumina sequencing; distance value of one indicating no community structure overlap, and a value of zero indicating identical community structure

Pearson correlation of the relative abundance of each taxon between Nanopore and Illumina data was conducted to further evaluate the reproducibility of sequencing results between the two platforms. Significant correlation was found at all phylogenetic levels from phylum to genus (R values > 0.98, p values < 0.001), indicating reproducible taxonomic assignment results can be obtained between Nanopore and Illumina platforms.

It should be noted in these comparisons that different primer sets were used for the two analysis approaches, which has previously been shown to bias microbiome community structure as analyzed by 16S rRNA sequencing (18). Despite this additional source of bias, the above analyses imply that the weighted community structure is comparable between the two platforms, encouraging future development.

### Between-Run Sample Carryover

We noted an apparent carryover of sequences between pure culture runs of *P. fluorescens* and *M. smegmatis*. The potential for sequence carryover has been anecdotally reported in the literature (19). We subsequently excluded all runs except the first run on each flow cell from earlier analyses, and undertook a formal evaluation of sequence carryover.

Results from analysis of the *Mycobacterium smegmatis* run are shown in Figure 4. In this run, 76.5% of sequences were assigned to the correct Actinobacteria phyla, 11.0% of sequences were incorrectly assigned to another domain or unable to be assigned, and 12.4% of sequences were incorrectly assigned to the Proteobacteria phyla, presumptively resulting from sequence carryover. 54.5% of sequences were assigned to the correct *Mycobacterium* genus whereas 10.8% of sequences were assigned to the *Pseudomonas* genus.

**Figure 4.**
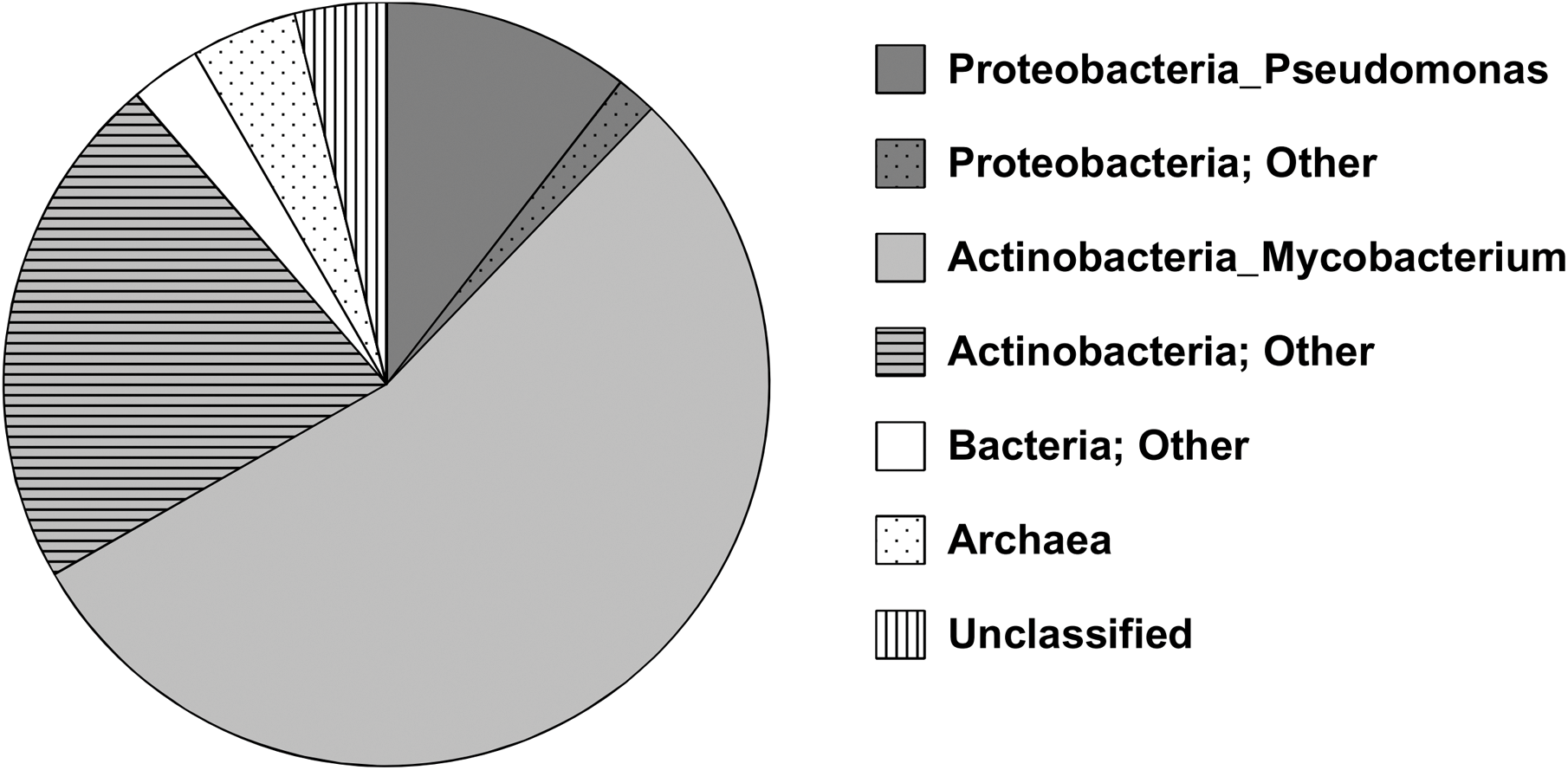
Taxonomy assignment of *Mycobacterium smegmatis* 16S rRNA pure culture sequencing following *Pseudomonas fluorescens* sequencing.

Following the *P. fluorescens* and *M. smegmatis* runs, we subsequently ran an additional environmental sample derived from river water and re-ran a *M. smegmatis* sample using 1D technology. In the second *M. smegmatis* run, we observed only 0.5% of sequences to be assigned to the correct Mycobacteriaceae family, compared with 54.5% in the first *M. smegmatis* run. These results imply that continued carry-over serves to significantly decrease output quality; however, additional validation with controlled microbial community composition is necessary to confirm this observation.

### Areas of Future Development

This investigation has identified multiple necessary areas of future development to enable 16S rRNA microbiome characterization on the Nanopore platform. First, strategies to exclude between-run carryover, either via improved washing between runs or a barcode approach, would enable multiple runs on the same flow cell, significantly reducing per-run costs. In this investigation we performed six runs on the same flow cell while observing minimal output loss. Second, improved bioinformatics strategies are necessary to exclude poor quality sequences. In the *E. coli* and *P. fluorescens* runs, 3.3 and 9.0% of sequences, respectively, were not assigned to any phyla, suggesting poor sequence quality and confounding both alpha- and beta-diversity analyses. Finally, it would be beneficial to develop a 16S rRNA annotation pipeline based upon optimized analysis strategies that provides output in near real-time, facilitating field and clinic applications and alleviating current bioinformatics challenges from interested investigators. Ultimately, the development of a rapid and low-cost microbiome approach will facilitate the application of clinical and environmental microbiome technologies.

## Materials and Methods

### Overview of DNA samples and sequencing libraries

Three pure culture bacterial DNA samples and an environmental DNA sample were analyzed in the current study. The three pure culture bacterial DNA samples were extracted from pure cultures of: (1) *Escherichia coli* (ATCC 15597), (2) *Pseudomonas fluorescens* (ATCC 13525), and (3) *Mycobacterium smegmatis* str. mc2 155, respectively. The environmental DNA samples, which had previously undergone 16S rRNA gene sequencing using the Illumina MiSeq platform, were a hydraulic fracturing produced water sample (17) and a river water sample (unpublished).

All Nanopore sequencing runs were conducted using the MinION Mk IB platform following recommended sequencing protocols (Oxford Nanopore Technologies). The *E. coli* 16S rRNA amplicon 2D library and the hydraulic fracturing produced water 16S rRNA amplicon 2D library were sequenced individually using a Nanopore MIN-105 flow cell and a Nanopore MIN-106 flow cell, respectively. The remaining libraries were sequenced on a MIN-106 flow cell following a sequential order: (1) *P. fluorescens* 2D library; (2) *M. smegmatis* 2D library; (3) river water sample 2D library; (4) *M. smegmatis* 1D library. For the sequential sequencing runs, flow cell washing was conducted immediately following the completion of the previous sequencing run using a Nanopore washing kit WSH002 (Oxford Nanopore). The Oxford Nanopore recommended washing protocol was used between runs, namely 150 µL of WSH002 solution A was loaded to the flow cell through priming port and incubated at room temperature for 10 minutes, then 150 µL of WSH002 solution B was loaded through the priming port before the next sequencing run and incubated for another 10 minutes at room temperature.

### Nanopore sequencing library preparation

Previously described universal primers targeting the 16S rRNA gene region (S-D-bact-0008-c-S20 and S-D-bact-1391-a-A-17) (20) were used for PCR. Each PCR was conducted in a total volume of 50 µL, containing 5 µL 10x buffer, 5 µL dNTP mix, 2.5 µL of each forward and reverse primer, 0.25 µL DreamTaq, 1 µL template DNA, and 33.75 uL nuclease free molecular grade water. The temperature condition for the PCR was 3 minutes at 95ºC; 30 cycles composed of 20 seconds at 95 ºC, 30 seconds at 47 ºC for annealing, 1 minute at 72 ºC; and a final elongation at 72 ºC for 15 minutes. All PCR products were purified using Ampure XP beads and normalized to 45 µL containing 1 µg of purified PCR products. Negative controls were used for all PCR reactions and DNA extractions, and all controls were negative.

2D libraries were prepared using a Nanopore NSK007 sequencing kit and recommended protocol (Oxford Nanopore Technologies). The end repair step of the purified PCR products was conducted by adding 7 µL Ultra II End-Prep buffer, 3 µL Ultra II End-Prep enzyme mix (New England Biolabs), and 5 µL control DNA provided with Nanopore NSK007 sequencing kit. The end repair reaction mix was incubated at 20ºC for 5 minutes and 65ºC for 5 minutes. The end-repaired PCR products were further purified using AMPure XP beads and ligated to the sequencing adapters by adding 8 µL molecular grade water, 10 µL Nanopore NSK007 adapter mix, 2 µL Nanopore NSK007 HPA solution, and 50 µL Blunt/TA Master Mix (New England Biolabs), and then incubated at room temperature for 10 minutes. 1 µL HPT solution from the NSK007 kit was added and incubated for an additional 10 minutes at room temperature. The ligated and tethered 2D libraries were purified by using MyOne C1 beads (Thermo Scientific) and eluted in 25 µL elution buffer (Oxford Nanopore Technologies). A description of the *M. smegmatis* 1D library preparation is included in the Supplementary Information.

All sequencing flow cells were primed using 500 µL Running Buffer Fuel Mix diluted in 500 µL molecular grade water following the recommended priming protocol (Oxford Nanopore Technologies). After priming, 6 µL of each 2D sequencing library was mixed with 37.5 µL Running Buffer Fuel Mix (Oxford Nanopore Technologies) and 31.5 µL molecular grade water, then loaded to Nanopore flow cell for sequencing.

## Sequence data processing

### Base-calling and initial format conversion

The raw FAST5 files were base-called using Metrichor v2.42.2 with 2D Basecalling for FLO-MIN106 250bps workflow and 1D Basecalling for FLO-MIN106 450bps workflow. Passed 2D reads of 2D sequencing libraries and passed template reads of 1D sequencing libraries were converted to FASTA files for downstream analysis using Poretools (21).

### Operational Taxonomic Unit Evaluation

Operational taxonomic unit (OTU) picking was conducted by using both closed-reference and *de novo* picking strategies implemented in QIIME 1.9.2 (22). Closed-reference OTU picking was conducted by using Greengenes 13.8 as the reference database (23).

### Taxonomy Assignment

For pure culture *E. coli* and *P. fluorescens* sequencing data, taxonomy was assigned to each sequence read within QIIME 1.9.0 (24) using the RDP classifier (25) against Greengenes 13.8 (23) and RDP 16S rRNA training set (25) as the reference database, respectively; as well as using BLASTn (26) against Greengenes 13.8 (23) as a reference database. For subsequent analyses, taxonomy was assigned to each individual sequence read using the RDP classifier (25) against Greengenes 13.8 (23) as this approach was found to achieve the highest taxonomy assignment accuracy for the pure culture *E. coli* and *P. fluorescens* sequence data.

Illumina 16S rRNA amplicon sequencing data of the produced water and river water samples were re-processed using the same approach as Nanopore sequence data. The Illumina data were clustered into OTUs using 100% similarity threshold, i.e. each identical Illumina sequencing read was assigned taxonomy using RDP classifier (25) against Greengenes 13.8 (23).

Jaccard and Bray-Curtis distances, which are dissimilarity distances measuring level of dissimilarity between two microbial communities, were calculated using QIIME 1.9.2 (22) to measure the degree of similarity of the produced water microbial community similarity between Nanopore and Illumina platforms. Significance of correlation between the two technical replicates of the produced water sample across sequencing platforms was conducted using Pearson’s correlation implemented in Minitab 16.

## Acknowledgements

Support for this project was provided by the University of Pittsburgh Central Research Development fund. KB was a member of the Oxford Nanopore Early Access Program, which initially provided access to the sequencing platform at reduced cost.

---

Table S1. Relative abundance distribution at phylum level for the produced water sample.

Table S2. Relative abundance distribution of Firmicutes phylum for the produced water sample.

